# Global epistasis in ecosystems arises from resource constraints

**DOI:** 10.64898/2026.05.12.724736

**Authors:** Seppe Kuehn

## Abstract

Global epistasis refers to the observation that the effect of a mutation or modification depends on the state of a biological system, not its detailed composition. Such patterns have been reported across biological scales, from proteins to organisms and ecosystems. In its simplest form, global epistasis appears as a linear relationship between the change in function or fitness due to a perturbation, and the background level of function or fitness. The mechanistic basis of global epistasis, particularly in ecological systems, remains unresolved. Here, we propose that in microbial communities, global epistasis describing the impact of adding a species to a community on function arises generically from constraints imposed by shared resource pools. We illustrate this mechanism in a single-species system growing on multiple substitutable resources, where global epistasis follows directly from nutrient limitation by an essential non-substitutable resource. We then extend this framework to multi-species communities competing for a single resource and show that the marginal effect of adding a species depends linearly on background community function, with a slope determined by the fraction of the resource claimed by the added species. We show that global epistasis persists in trophic cascades, but that facilitation and niche partitioning qualitatively break the linear dependence. This study provides a simple explanation for the appearance of global epistasis in ecosystems, and suggests that global epistasis should be a null expectation in ecosystems governed by competition. Our results propose that coupling between perturbations and shared resource pools might also help explain global epistasis at the organismal level.

## I. INTRODUCTION

Fitness landscapes describe the impact of genotypic variation on the fitness of organisms or the function of enzymes. Landscapes characterize the additive and epistatic impacts of the presence and absence of mutations in genes or genomes on activity or fitness[1]. Landscapes provide a quantitative statistical lens for understanding how changes in the makeup of a biological system impact a functional property of interest.

Recently, the idea of landscapes has been extended to ecosystems, where they describe not fitness but ecosystem function. In this context, community composition takes on the role of genotype. Thus, community-function landscapes [2] describe how changes in community composition impact functional properties from biomass productivity [3], to compound degradation [4], and pathogen inhibition [5].

Studies have demonstrated the existence of simple patterns that capture epistatic properties of landscapes in proteins [6], organismal fitness [7, 8], and community function [9]. These patterns propose that there are global constraints on the underlying structure of landscapes that render fitness or function more predictable than one would naively expect. A primary example of this pattern is global epistasis, where the effect of a genotypic change on fitness or function depends not on the specific background (e.g., which mutations or species are present in a protein or ecosystem) but on the system’s baseline function. In the simplest, and most widely observed case, global epistasis manifests as a decreasing *linear* dependence of the impact of a mutation or species addition on background enzyme or ecosystem function. For example, the effect of mutations on yeast fitness are observed to depend linearly on the fitness of the organism in the absence of the mutation [7]. In this case, mechanisms for global epistasis have been proposed, including statistical effects and cellular resource constraints [10, 11], but not yet validated empirically.

Here we focus on such linear patterns of global epistasis in the context of microbial ecosystems [9] where adding a species to a microbial community modifies the function in a manner that depends only linearly on the function of the community in the absence of the added species (Figure 1). Global epistasis in microbial communities is observed for functions such as the total biomass, production or degradation of a target compound, or inhibition of a pathogen, results that extend to plant communities. In these communities the origins of these linear dependencies remain unclear. Here, we propose the idea that these patterns emerge generically from systems where there is competition for a shared (global) resource that all components of the system must compete for. In communities, this resource is typically a nutrient or shared substrate required for growth.

**FIG. 1.**
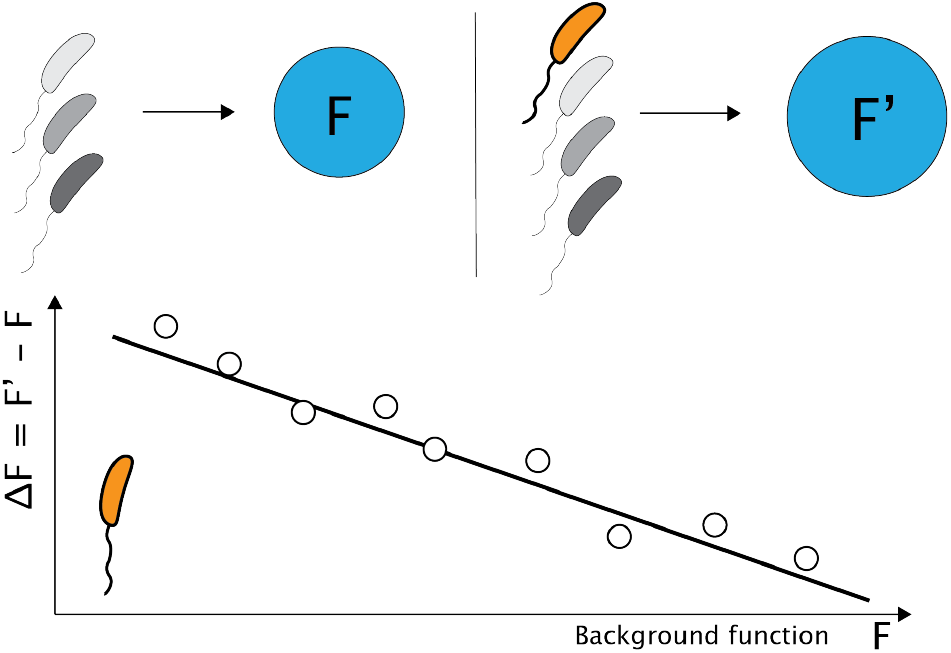
Global epistasis in microbial communities. Conceptual illustration of global epistasis in microbial communities. Consider a background community (gray cells) with some function (*F*) which could be compound production, biomass, or pathogen inhibition. Then consider the addition of a single target strain (orange) to this background community resulting in a new function (*F*′). Lower plot shows the change in function (Δ*F* = *F*′ – *F*) as a function of the background community function *F* . Global epistasis is the linear dependence of Δ*F* on *F* .

## II. RESULTS

Recent studies have shown that the notion of a landscape can be generalized to understanding microbial community function, such as compound production or degradation. Landscapes defining community function map the composition of the community (species presence/absence) to a functional property of interest. In this context, global epistasis as discussed above manifests as a linear dependence of a change in community function (Δ*F*) upon the addition of a species to a background community on the background community function (*F*).

Figure 1 demonstrates the basic idea. A community utilizes some resource pool such as carbon, nitrogen, or light. Function (*F*) is then the product of growth on that resource which could be total biomass production, conversion of the resource to a specific compound such as butyrate in the gut microbiome, or the degradation of a catabolite (blue circles). For low functioning communities (upper left, Figure 1), the addition of a new species (orange) results in a large increase in community function. In contrast, a high functioning community can only be marginally improved by the addition of the same species. The linear dependence of Δ*F* on *F* is defined as global epistasis. The goal of the present study is to show that this dependence emerges naturally in microbial communities from competition for a shared resource.

### Global epistasis in biomass production of one species on many resources

We begin with a simple example to illustrate how resource constraints can give rise to global epistasis. In this scenario the relevant landscape is the growth of a single strain on multiple substitutable resources, such as a microbe utilizing an array of carbon sources. The situation is depicted in Figure 2a-b. Function is defined as the total biomass produced by a single species. Crucially, growth also depends on another non-substitutable resource (nitrogen, star - green, Figure 2a) which is present in a fixed quantity.

**FIG. 2.**
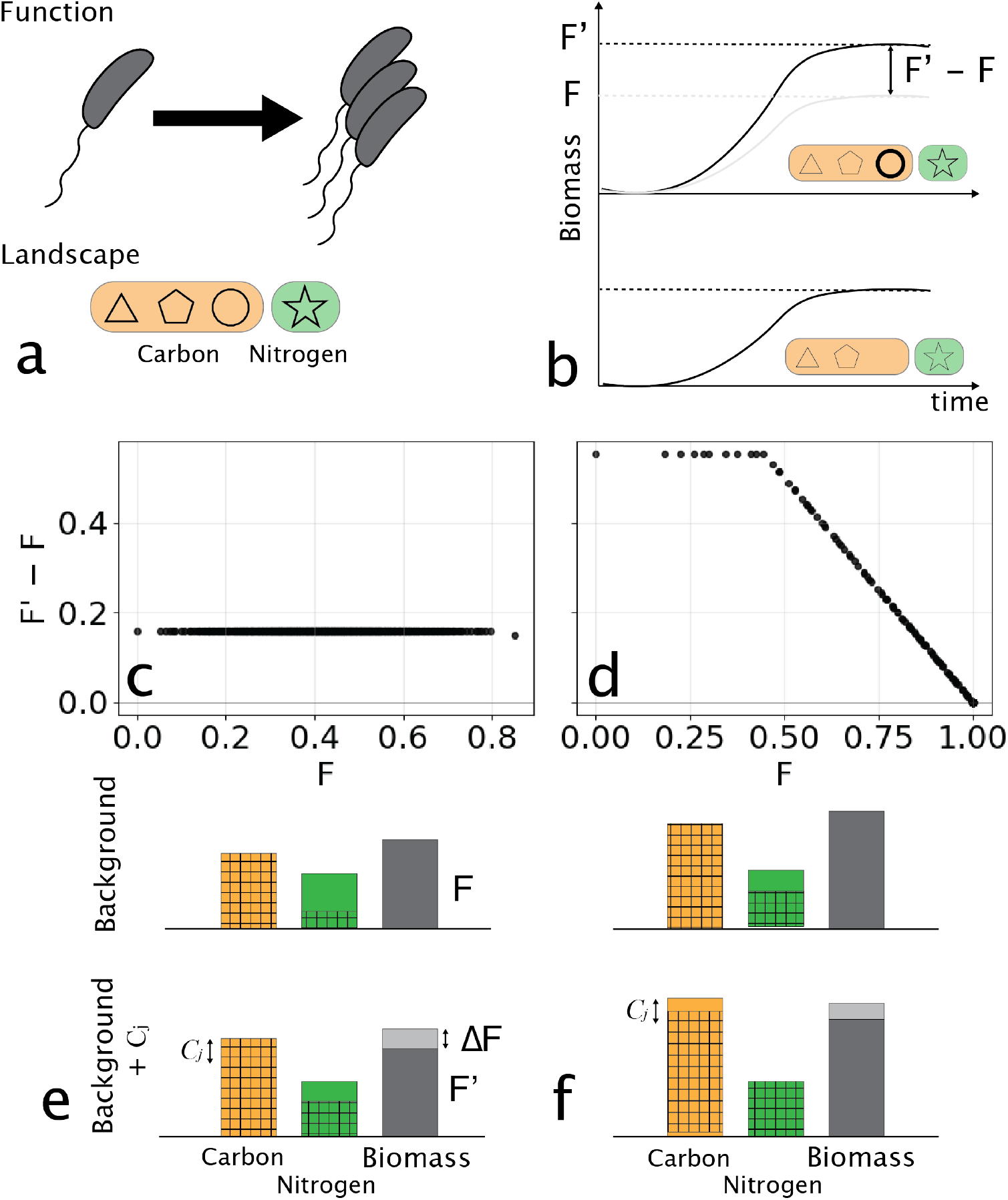
Global epistasis emerges from simple resource budgets in a one–species, many–carbon source model. **(a–b)** Conceptual overview. A single microbial population consumes multiple substitutable carbon sources (orange, shapes) and one essential nitrogen source (green). Shapes denote distinct carbon sources. Panel (a) illustrates the resource pool and a background set of available carbons; panel (b) shows how adding a focal carbon source *C*_*j*_ (bold circle) increases total biomass growth *F* when nitrogen is in excess. **(c–d)** Global epistasis plots for a focal carbon source *C*_*j*_ across all 2^9^ background combinations of the remaining carbon sources. Each point compares the biomass produced in a background *S* (*F*, x-axis) with the additional biomass gained by adding *C*_*j*_ (*F*′ – *F*, y-axis). *(c) Carbon-limited backgrounds:* adding *C*_*j*_ increases the carbon budget by a fixed amount 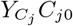, producing an approximately constant biomass gain and a horizontal line. *(d) Nitrogen-limited backgrounds:* as *F* approaches the nitrogen budget, the gain from adding *C*_*j*_ is limited by the remaining nitrogen, yielding a linear decline in *F*′ – *F* with a slope –1. **(e–f)** Summary of resource usage and biomass yields across representative background conditions for the cases shown in (c) and (d) respectively. Orange bars show total available carbon, green bars show nitrogen, and gray bars indicate total biomass production. Top row illustrates background resource usage without the target resource added in a carbon limited (e) and nitrogen limited (f) scenario. Bar heights are resource availability, and hatched areas resource utilized. Bottom row indicates resource utilization with the target resource added. Gray portion of the biomass bar indicates added biomass upon addition of *C*_*j*_ (Δ*F*). Note that nitrogen is never limiting in (e), but becomes limiting with the addition of *C*_*j*_ in (f).

The consumer resource model for a single strain growing on a set of substitutable carbon sources and a single non-substitutable nitrogen source is

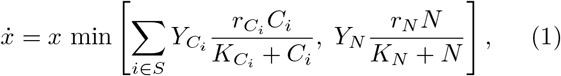

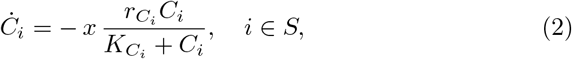

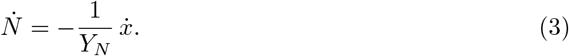

Growth of the two non-substitutable nutrients is governed by Leibig’s law of the minimum. *r*_*_ represent rates of uptake, *Y*_*_ biomass yields, and *K*_*_ affinity parameters. The landscape is defined by the distinct carbon sources available for growth, and function in simply *x*(*t* → ∞). The change in biomass (function, Δ*F*) due to the addition of a carbon source (*C*_*j*_) is then determined by integration of the model until growth stops and comparing the final biomass to that without the target carbon source added (Figure 2b).

Figure 2c-d shows the two main results from this simple model. Here we consider a landscape defined by the availability of 10 distinct resources (*C*_*j*_) which vary in the corresponding growth yields (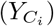) and uptake rates (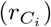). In panel c the nitrogen resource is in excess, and the function is purely determined by the carbon availability. A single target *C*_*j*_ is held out, and for all possible combinations of the remaining 9 resources, the growth is computed (*F*). We then compute the additional growth upon inclusion of *C*_*j*_ to each background resource combination, and observe a flat line with intercept 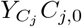. This flat line reflects the constant additional biomass produced upon the addition of this carbon source. Note that nitrogen is never limiting. A schematic illustration of resource utilization in this scenario is shown in Figure 2e.

Contrast the example in Figure 2c with the case where nitrogen limitation occurs upon the addition of the target carbon source in panels (d,f). In this case, nearly all of the nitrogen is exhausted during growth on the background carbon sources, and the addition of the target carbon source causes nitrogen to be completely utilized while some carbon is retained in the system as shown schematically in Figure 2f. In this case, we see the hallmark of global epistasis, which is a linear relationship between the background function (F, Figure 2d) and the change in function (*F*′ – *F*), here arising from nutrient limitation.

In the appendix, we show that this decline in Δ*F* due to nutrient limitation gives rise to a slope of –1. Furthermore, Δ*F* is strictly positive, and cannot cross over zero, which occurs when nitrogen is fully consumed in the background nutrient condition. This arises simply because adding nutrients in this model cannot reduce biomass production. Nonetheless, this example serves to demonstrate the route to apparent global epistasis, in this case for a landscape defined by nutrient availability, due to resource limitation. Next we turn our attention to a situation where epistasis arises in a landscape defined by species presence/absence rather than resource presence/absence.

### Global epistasis in competitive communities

The previous example serves to demonstrate a simple case where resource limitation can give rise to patterns that evoke global epistasis. However, this first example has three limitations. First, in theory it only permits Δ*F* vs *F* slopes of –1, while these slopes are observed to vary [9]. Second, Δ*F* is often observed to be negative, an outcome which is not possible in the previous example. Third, global epistasis in communities is observed in the impact of species presence/absence on the communities, not resources. Here, we address these limitations in the context of competitive communities, again finding a central role for resource limitation and competition.

Consider a community with *N* species where each competes for a central resource and excretes a byproduct, such as a fermentation product. In this situation, species are defined by the parameters that control their growth on the central resource (rate, yield) and the yield of byproduct during growth. A consumer-resource model of this scenario is:

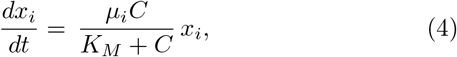

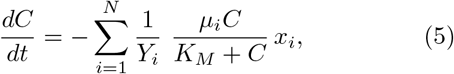

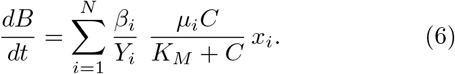

*x*_*i*_, *C* and *B* are the biomass of strain *i*, a carbon source, and a byproduct respectively. *µ*_*i*_ is the growth rate of strain *i* on *C, β*_*i*_ is a yield of byproduct per unit carbon consumed for strain *i* and *Y*_*i*_ the growth yield on the carbon substrate. Phenotypes of individual taxa are determined by the parameters that govern growth and excretion (*µ*_*i*_, *β*_*i*_, and *Y*_*i*_), and the landscape is defined by strain presence and absence.

To construct a global epistasis plot in this context we simulate a community of *S* strains until growth stops (*C* → 0) and compute the total accumulated byproduct (*F* = *B*(*t* → ∞)). Repeat this process for the same community with the target species added and compute *F*′ (Figure 3a,b). This process is carried out for all possible background communities.

**FIG. 3.**
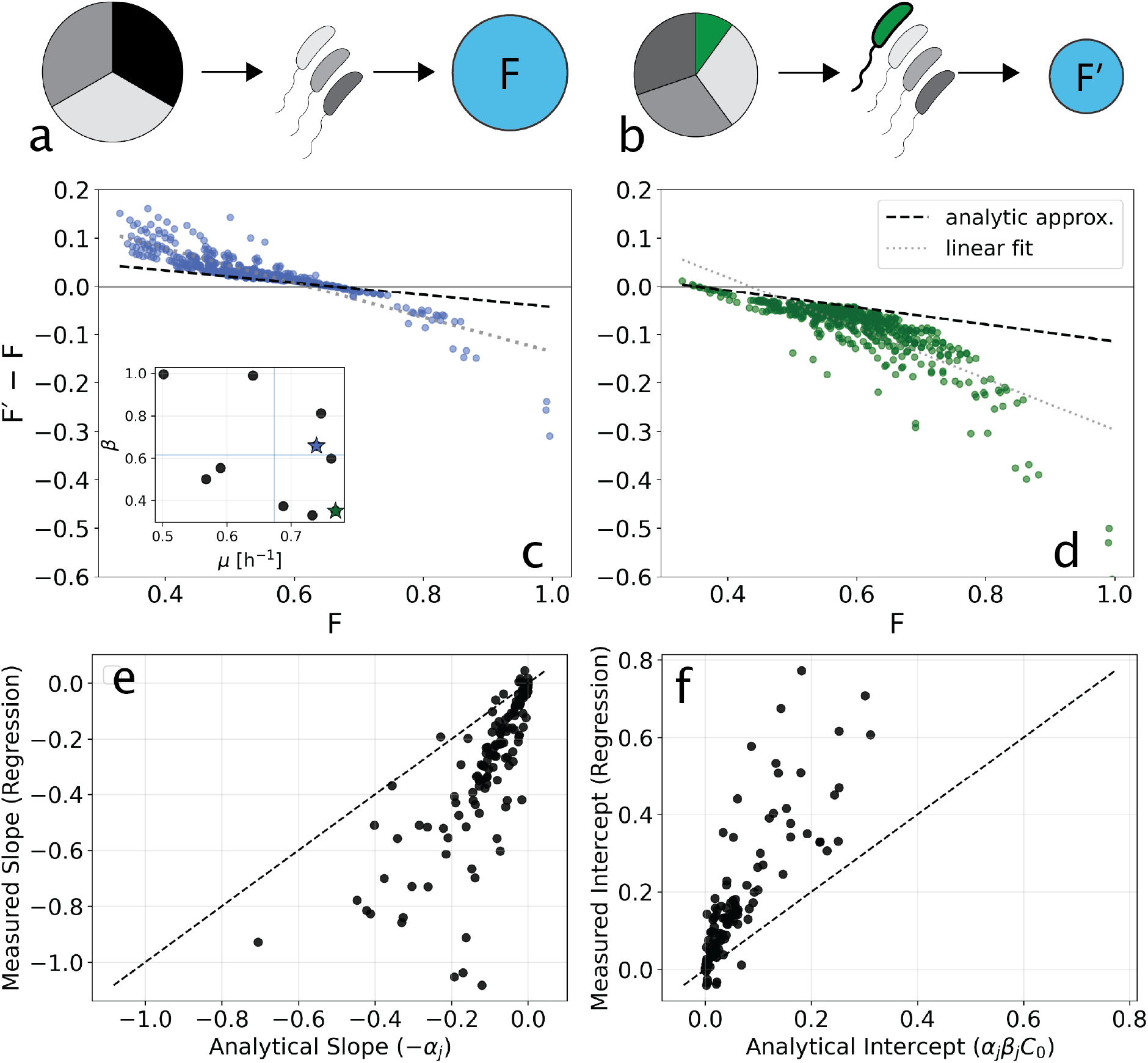
Global epistasis in competitive communities. Simulations for the model shown in Eqns. 4-Δ **(a)** A schematic of a background community consuming the supplied carbon in equal proportions across all three members. **(b)** An added member (green) consumes a portion of the supplied carbon (green wedge), but the remaining resource is consumed in roughly equal proportions by the background community. **(c,d)** Global epistasis plots of the change in community function against background function. Simulations for a community defined by 10 species which vary depending on their maximum growth rates (*µ*_*i*_) and byproduct production per carbon consumed (*β*_*i*_, y-axis) inset, panel (c). Each point corresponds to a strain, colored stars correspond to target species in (c, d). Heavy dashed lines are the analytical approximation in Eqn. 7 and dotted lines are linear regressions. **(e,f)** Compare analytical and measured slopes (e) and intercepts (f) of global epistasis plots like those shown in (c-d) 14 random 10-species communities. The dashed line indicates equivalence between measured and analytical predictions.

A target species that grows slowly relative to the background community will have little impact on function, since it consumes little of the primary resource due to lagging other members of the community. The result will be a horizontal line near zero since *F*′ – *F ≈* 0 (examples in Suppl. Figure S1).

In contrast, if the target species is competitive for the carbon source due to a high growth rate, it will impact community function. Two such examples are shown in Figure 3c,d. The growth rates *µ*_*i*_ and byproduct production rates *β*_*i*_ are shown in the inset to (c) for all species. For a species that has moderate production of byproduct (e.g. blue star, Figure 3c inset) and relatively fast growth rate it consumes a larger fraction of the available carbon source when added to the background community. In this scenario the target species can increase community function relative to the background for small *F* because a larger fraction of the carbon is consumed by a moderate producer of byproduct. At high background function, the impact becomes negative due to fast growth and only moderate *β*_*i*_ of the target species reducing overall byproduct production. The result is *F*′ – *F <* 0. For a species of high growth rate and low byproduct production, Δ*F <* 0 for almost all backgrounds due to fast growth consuming significant primary resource without substantial byproduct production (Figure 3c inset, and panel d). Despite these intuitive observations, it is not obvious why this model should give rise to an approximate linear dependence between Δ*F* and *F* . We investigated this analytically, and a detailed calculation is shown in the appendix. Here we recapitulate the essential features of this calculation.

### Linear dependence of Δ*F* **on** *F*

To gain an intuition for why we observe approximately linear changes in Δ*F* with *F* first consider a background community with composition *S* which does not include species *j*. The community *S*^′^ = *S j* is the community with *j* added. So Δ*F* = *F* (*S*^′^) *F* (*S*).

As the community *S* consumes the initial carbon (*C*(0)), they produce a total by-product, *F* . This output is determined by the community’s average efficiency, ⟨*β*⟩_*S*_, such that *F* = *C*_0_⟨*β*⟩_*S*_. When we re-run growth with the target species added (*j*) it claims a fraction of the total carbon pie during growth with the background community. Define this fraction as *α*_*j*_ (unitless). The net change in by-product, Δ*F* = *F*^′^ – *F*, is simply the difference between the new production contributed by species *j* and the production lost from the existing community because their carbon share has been reduced.

Species *j* produces an amount of by-product equal to its share of the initial carbon multiplied by its own efficiency or *α*_*j*_*C*_0_*β*_*j*_. At the same time, the existing community loses a portion of its resources equal to *α*_*j*_*C*_0_. Since the community was converting carbon into by-product with an average efficiency of ⟨*β*⟩_*S*_, the amount of by-product lost is (*α*_*j*_*C*_0_) ⟨*β*⟩_*S*_. Because the average efficiency ⟨*β*⟩_*S*_ is just the total by-product *F* divided by the total carbon *C*_0_, this loss simplifies to *α*_*j*_*F* .

By combining these two terms, the gain in byproduct from species *j* and the lost byproduct due to carbon utilized by *j* we find that the net change is

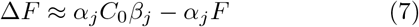

which is linear in *F* . This result intuitively explains why we see straight lines in global epistasis experiments: the gain in function from the new species is a constant, while the loss from the background is strictly proportional to the initial productivity *F* . As the background becomes more productive, the proportional loss grows larger, causing the net benefit of adding the new species to decline at a constant rate defined by the slope *α*_*j*_. This approximation is shown as the heavy dashed lines in Figure 3c, d.

The linearity depends on an assumption of invariant internal allocation: the requirement that the addition of species *j* does not alter the relative proportions of carbon consumed among the background species. This assumption will clearly break down if there is significant variance in growth rates (*µ*_*i*_) or yields (*Y*_*i*_) across the community. For example, adding a highly efficient competitor *j* reduces the total time available for growth by more rapidly exhausting the common resource. This disproportionately penalizes slow-growing species compared to fast-growing ones, causing the background efficiency ⟨*β*⟩_*S*_ to shift during the transition from *S* to *S*^′^. However, as we will see empirically, even when this assumption breaks down, global epistasis plots remain approximately linear.

In Figure 3c,d we see evidence of this type of deviation from the analytical approximation at large values of *F* . In this scenario, adding the relatively fast growing target species will diminish the impact of slow growers, and if those slow growers are significant producers of byproduct this will penalize community function beyond the analytic approximation (points at high *F* below the heavy dashed line). However, even with these departures from our analytical expectations, we note that in the plots shown in Figure 3(c, d) the data follow an approximate linear relationship. The light dotted lines show linear regressions which exhibit a larger slope and higher intercept than the analytical approximation as expected.

To test the analytical approximation more broadly, we simulated many random 10 species communities with increasing interspecies variability in the growth rates (see Suppl. Fig. S2). We computed global epistasis plots for all members of each community and plotted the measured (regression) slopes and intercepts against our analytical expectations (Figure 3e,f). As expected, we find that regression slopes are larger (more negative) and intercepts higher than analytical approximations. These deviations arise from the departures from our analytical assumptions described above.

Next we wanted to understand how more complex ecological interactions might complicate the picture of global epistasis arising from a resource constraint.

### Global epistasis in the presence of cross-feeding

While competitive interactions are known to be important in microbiomes [12], many communities are structured in metabolic cascades. In this context, one competitive group produces an intermediate compound that is then used as input to the next trophic layer. Examples include fermentation cascades in bioreactors [13], or denitrification cascades in soils [14]. Here, we investigate whether the approximately linear dependence of Δ*F* on *F* is retained when communities are structured trophically.

We extended the consumer resource model from the previous section to include a primary carbon source that is consumed by producer strains that excrete a second carbon source (*C*_*cf*_, Figure 4a) [15]. This introduces two modifications relative to the previous model. First, there are now two types of strains: producers (*x*_*p*_) that utilize the carbon source *C* and produce a secondary metabolite *C*_*cf*_ in proportion (*p*_*p*_) to the carbon consumed, and consumers (*x*_*k*_) that then utilize *C*_*cf*_ . Both types produce byproduct *B* and, as before, function is accumulated *B*. The full system of ODEs is given in the SI (Eq. (25)).

**FIG. 4.**
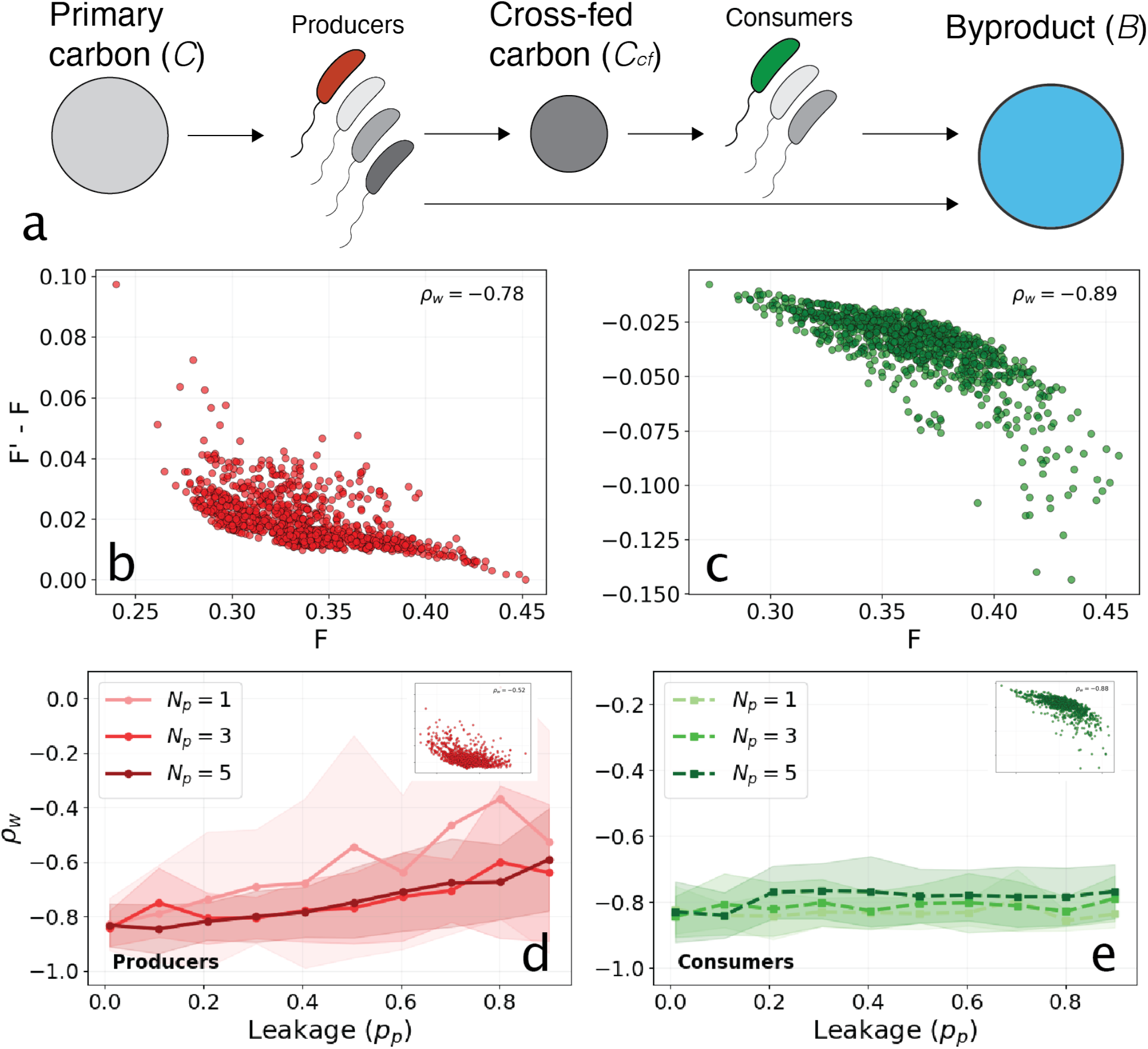
Global epistasis in the presence of cross feeding. (a) Schematic of the metabolic cascade: producers take up primary carbon (*C*) and release cross-fed byproduct (*C*_*cf*_, with fraction *p*_*p*_), which fuels the consumer guild. All species contribute to a final byproduct pool (*B*, via coefficient *β*\*). (b, c) Example global epistasis plots showing function gain (*F*′ – *F*) against background function (*F*) in the low-leakage regime (*p*_*p*_ = 0.1), for a focal target species that is a producer (b, red) and a consumer (c, green). Each scatter cloud represents one focal target across many background communities. (d, e) Density-weighted correlation coefficient *ρ*_*w*_ of Δ*F* vs *F* as a function of leakage (*p*_*p*_), averaged across all producer target species (d) and all consumer target species (e) within each community, and then averaged across 10 random community parameterizations (*N* = 15 species). Shaded regions represent 1 standard deviation across the 10 randomizations. Lines correspond to communities with different numbers of producer species (*N*_*p*_) as shown in the legend. Increasing *ρ*_*w*_ (less negative) in (d) is indicative of a breakdown of the linear GE regime, which is not observed for the consumer guild (e). Insets show example global epistasis plots at *p*_*p*_ = 0.8 for a focal producer (d, inset) and a focal consumer (e, inset).

Simulating this model reveals approximately linear relationships between Δ*F* and *F* for both guilds. Figure 4b and 4c show global epistasis plots for a focal target species that is a producer (b) and a consumer (c), respectively, at low leakage (*p*_*p*_ = 0.1). In each case the scatter is taken across many background communities, and despite some curvature the density-weighted correlation coefficients (*ρ*_*w*_) approach –1. In this low-leakage scenario the production of byproduct is dominated by the producers and the dynamics are similar to the purely competitive community above.

To investigate the robustness of this pattern we simulated communities with varying numbers of producers (*N*_*p*_) across a range of leakage values. For each community we computed *ρ*_*w*_ for every member of each group and averaged the result within that group. Figure 4d shows this group-averaged *ρ*_*w*_ as a function of leakage for the producers, that is, the average across global epistasis plots in which the target species is a producer. Figure 4e shows the same where the target species is a consumer. As the fraction of primary carbon that producers leak into the cross-fed pool rises, the producer-group *ρ*_*w*_ becomes less negative, signaling a departure from approximately linear behavior, while the consumer-group *ρ*_*w*_ remains close to –1 throughout.

The departure from global linearity observed at high leakage (*p*_*p*_ = 0.8, Figure 4d, inset) reflects saturation of the cross-fed metabolite pool. The magnitude of Δ*f* is largest when the background community contains few producers and therefore generates little *C*_*cf*_, so the focal producer’s contribution substantially augments the resource available to the consumer group. As the number of producers in the background rises, the existing producers drive *C*_*cf*_ toward the ceiling set by the supplied primary carbon (*C*_0_), and each additional producer has diminishing effect. In this saturated limit Δ*F* flattens, producing the curved Δ*F* vs *F* relationship in the inset and the corresponding decline of the weighted correlation coefficient to *ρ*_*w*_ *≈* –0.5.

In contrast to the producer group, global epistasis plots for focal consumer target species maintain high linearity (*ρ*_*w*_ *≈* 0.9, Figure 4c,e) even at elevated leakage levels. This persistence arises because consumers act as secondary exploiters of a resource pool (*C*_*cf*_) that is bounded by the primary intake of the producers, this results in competition for an approximately fixed *C*_*cf*_ . Consequently, function gains remain inversely proportional to the total community function via the competitive mechanism from the previous section.

The preceding examples reveal that an approximately linear dependence between Δ*F* and *F* should be expected in communities that include trophic structure.

### Niche partitioning and facilitation break global epistasis patterns

The above examples, competition, and cross feeding show approximately linear relationships between Δ*F* and *F* . As one would expect, there are scenarios where these linear trends can be broken. Here we briefly discuss two such scenarios. A complete treatment of these two cases is shown in the Appendix.

#### Niche partitioning

First is the case where there are two independent groups consuming non-overlapping resources [16]. These might be guilds or metabolic groups that are not competing for any shared resource. In this scenario there are competitive interactions between members of the same group, giving linear trends similar to Figure 3. However, when the target species is held out from a background composed of only members of the other group there is no effect. The result is a Δ*F* vs. *F* plot that has distinct regimes (linear declining and flat) giving rise to no clear linear trend across all backgrounds (Figure S4a).

#### Facilitation

A second example is facilitation [17]. In this case one strain alters the behavior of another in a manner favorable to function. Concretely, if *β*_*i*_ is increased in a manner that depends on the presence of species *j* then the Δ*F* with *F* plot can exhibit positive slopes. An example of this behavior is shown in Figure S4b.

## III. DISCUSSION

Changes in microbial community function upon the addition of a new species exhibit decreasing linear trends with background community function due to the presence of shared resources. These observations are borne out in single species growing on multiple resources, multiple species competing for a centralized resource, and trophic cascades. The central contribution of this work is elucidating a simple, and plausible, ecological mechanism for these patterns.

This study suggests that global epistasis in communities arises from comparatively simple interactions mediated via a common resource pool that map to community function. Thus, in many metabolic contexts we might naively expect approximately linear relationships between Δ*F* and *F* . In this case, global epistasis might be considered the null expectation and qualitative departures from this behavior could be used as indicators of non-trivial ecological interactions impacting function. A concrete example of this is the positive slopes observed for a single strain in starch degrading communities [18].

Recent work in synthetic ecology proposes navigating landscapes that are defined in terms of species composition [2, 3]. In this view, the natural variables for understanding community function are compositional. However, as community richness rises, the dimensionality of compositional landscapes expands exponentially. Our results suggest that community function might be more naturally understood through the lens of resource dynamics than through community composition. Our results suggest that this complexity disguises a simpler underlying dynamics at the level of resources. Specifically, in the competitive communities discussed above, it is the average conversion of *C* to *B* by the background community and the growth and byproduct production of the target species that define *F* and Δ*F* . Thus, changes in the function of the community arise from changes in the efficiency of conversion of resources to product on average and on the impact that the target strain has on resource availability to the background community.

The idea that resource fluxes might be simpler variables for describing communities than abundances has been observed in other contexts. For example, communities often exhibit functional similarity despite substantial compositional variation [19], sometimes called functional redundancy [20], although clean null models for testing this idea quantitatively have only recently been put forward [21]. Similarly, recent work suggests that even in very diverse soil communities, metabolite dynamics can be described by models of a few effective biomasses [14] that coarse-grain the metabolic activity of hundreds of distinct taxa. In these contexts, it is the flux of metabolites through the system that provides access to a simple and interpretable phenomenology of community function.

Finally, global epistasis is widely observed in the context of the impact of mutations on fitness in evolving populations of microbes. There are several explanations for linear dependence of Δ*F* on *F* in this context, including statistical effects [10]. Our work suggests that such patterns might emerge from a coupling of mutational effects via global intracellular resources, such as ribosomes or metabolic enzymes. The proposal is supported by low-dimensional models of microbial growth that rely on the balance of protein synthesis and the production of charged tRNA [22]. Recent studies on the origins of global epistasis in organismal fitness have begun to explore this possibility [11]. As a result, global epistasis might arise from the constraints of sub-cellular resource pools and the statistical properties of mutations coupling to those constraints.

## ACKNOWLEDGMENTS

S.K. acknowledges conversations with Alvaro Sanchez and Mikhail Tikhonov. S.K. acknowledges the National Institute of General Medical Sciences R01GM151538, the National Science Foundation through the Center for Living Systems (grant no. 2317138), the National Institute for Mathematics and Theory in Biology (Simons Foundation award MP-TMPS-00005320 and National Science Foundation award DMS-2235451), and support from Department of Defense, ARO AWD106455.

## Notes

### Competing Interest Statement

The authors have declared no competing interest.

