## Supplement for "Global epistasis in ecosystems arises from resource constraints"

### SUPPLEMENTARY APPENDIX

#### GLOBAL EPISTASIS FROM ONE SPECIES MANY RESOURCES

Let $x(t)$ denote biomass; $C_i(t)$ the concentrations of carbon sources $i \in S$; and $N(t)$ the nitrogen concentration. Each carbon source is substitutable with uptake coefficient $r_{C_i}$, half–saturation $K_{C_i}$, and yield $Y_{C_i}$. Nitrogen is essential with $r_N$, $K_N$, and yield $Y_N$.

Under Liebig's law of the minimum, the instantaneous growth rate is

$$\dot{x} = x \,\min\left[\sum_{i\in S} Y_{C_i}\,\frac{r_{C_i}C_i}{K_{C_i}+C_i}\;,\; Y_N\frac{r_N N}{K_N+N}\right]. \tag{8}$$

Carbon and nitrogen dynamics follow from stoichiometry:

$$\dot{C}_i = -\,x\,\frac{r_{C_i}C_i}{K_{C_i}+C_i}, \qquad i\in S, \tag{9}$$

$$\dot{N} = -\frac{1}{Y_N}\,\dot{x}. \tag{10}$$

At batch termination, the total biomass gain is

$$F(S) = x(t_{\text{stop}}) - x(0).$$

##### Resource budgets

Let $C_{i0}$ and $N_0$ be initial stocks. Equation (8) implies the total biomass yield is limited by whichever budget can be fully consumed:

$$F(S) = \min\left(\sum_{i\in S} Y_{C_i}C_{i0},\; Y_N N_0\right). \tag{11}$$

Define the nitrogen budget

$$X_{\text{bud}} = Y_N N_0.$$

Thus:

- *C-limited:* $\sum_{i\in S} Y_{C_i}C_{i0} < X_{\text{bud}}$.
- *N-limited:* $\sum_{i\in S} Y_{C_i}C_{i0} \geq X_{\text{bud}}$.

##### Leave-one-in ($S' = S\cup\{j\}$)

*a. C-limited on both $S$ and $S'$.*

$$F(S') = F(S) + Y_{C_j}C_{j0}, \tag{12}$$

$$F' - F = Y_{C_j}C_{j0}. \tag{13}$$

All points lie on a horizontal line.

*b. N-limited on both.*

$$F(S) = F(S') = X_{\text{bud}} \;\Rightarrow\; F' - F = 0.$$

*c. Mixed regime.* If $S$ is C-limited but $S'$ crosses into N-limited:

$$F' - F = \min\big(Y_{C_j}C_{j0},\; X_{\text{bud}} - F\big)\,. \tag{14}$$

Equation (14) is piecewise linear: a segment of slope $-1$ as $F \to X_{\text{bud}}$, capped by the intrinsic contribution $Y_{C_j}C_{j0}$.

##### Interpretation

When nitrogen ultimately limits growth, adding $C_j$ can supply at most the remaining budget $X_{\text{bud}} - F$. This shrinks linearly with $F$, producing the characteristic straight lines with slope $-1$ near the N limit and horizontal plateaus elsewhere.

A compact summary is:

$$F' - F = \begin{cases} Y_{C_j}C_{j0}, & \text{C-limited } S, S', \\ 0, & \text{N-limited } S, S', \\ \min\{Y_{C_j}C_{j0},\; X_{\text{bud}} - F\}, & \text{mixed.} \end{cases}$$

#### GLOBAL EPISTASIS FROM MANY SPECIES, ONE RESOURCE

##### 1. MODEL DEFINITION

We consider a well-mixed batch culture containing $N$ microbial species that all compete for a single limiting carbon resource $C$ and produce a secreted by-product $B$ in proportion to their resource uptake. The dynamical variables are $C(t)$ (resource concentration), $B(t)$ (by-product concentration), and $x_i(t)$ (biomass of species $i$).

Each species $i$ is characterized by four parameters: $\mu_i$ (maximum specific growth rate), $K_M$ (shared Monod constant), $Y_i$ (biomass yield on carbon), and $\beta_i$ (by-product stoichiometry). The system evolves according to:

$$\frac{dx_i}{dt} = \mu_i\,\frac{C}{K_M + C}\,x_i \tag{15}$$

$$\frac{dC}{dt} = -\sum_{i=1}^{N}\frac{1}{Y_i}\,\mu_i\,\frac{C}{K_M + C}\,x_i \tag{16}$$

$$\frac{dB}{dt} = \sum_{i=1}^{N}\beta_i\,\frac{1}{Y_i}\,\mu_i\,\frac{C}{K_M + C}\,x_i \tag{17}$$

Growth ceases when the resource is depleted, $C(t_{\text{stop}}) \approx 0$. The quantity of interest is the final by-product $F(S) = B(t_{\text{stop}}; S)$, where $S$ denotes the subset of species present at inoculation.

#### 2. CHANGE OF VARIABLES AND SIMPLIFICATION

Since the Monod factor $C/(K_M + C)$ multiplies every species' growth rate, the dynamics simplify under the reparameterization $d\tau \equiv \frac{C}{K_M+C}\,dt$. Using this substitution, Eqs. (15)–(17) become:

$$\frac{dx_i}{d\tau} = \mu_i x_i, \quad \frac{dC}{d\tau} = -\sum_i \frac{1}{Y_i}\mu_i x_i, \quad \frac{dB}{d\tau} = \sum_i \beta_i \frac{1}{Y_i}\mu_i x_i. \tag{18}$$

These integrate exactly to $x_i(\tau) = x_{i0}\,e^{\mu_i \tau}$. Let $\tau_f$ be the effective time at carbon exhaustion ($C(\tau_f) = 0$). Integrating $dC/d\tau$ gives the carbon balance:

$$C_0 = \int_0^{\tau_f} \sum_{i\in S} \frac{1}{Y_i}\mu_i x_i(\tau)\,d\tau = \sum_{i\in S} \frac{x_{i0}}{Y_i}\left(e^{\mu_i \tau_f(S)} - 1\right). \tag{19}$$

We define $U_i(S) \equiv \frac{x_{i0}}{Y_i}\left(e^{\mu_i\tau_f(S)} - 1\right)$ as the total carbon consumed by species $i$, where $\sum_{i\in S} U_i(S) = C_0$. The final by-product is:

$$F(S) = \sum_{i\in S} \beta_i\, U_i(S). \tag{20}$$

#### 3. TARGET SPECIES ADDITION

Consider a held-out species $j$ and a background $S$. Define the augmented community $S' = S \cup \{j\}$. We analyze the change in by-product $\Delta F = F(S') - F(S)$.

##### 3.1. Decomposition of the Mean Yield

Define the carbon share of the added species $j$ as $\alpha_j = U_j(S')/C_0$. The remaining carbon is consumed by the background species $S$, such that $\sum_{k\in S} U_k(S') = C_0(1 - \alpha_j)$. We decompose the mean yield of the augmented community, $\langle\beta\rangle_{S'} = F(S')/C_0$, by splitting the sum:

$$\begin{aligned}
\langle\beta\rangle_{S'} &= \frac{1}{C_0}\left[\sum_{i\in S} \beta_i U_i(S') + \beta_j U_j(S')\right] \\
&= \frac{\sum_{k\in S} U_k(S')}{C_0}\left(\frac{\sum_{i\in S}\beta_i U_i(S')}{\sum_{k\in S} U_k(S')}\right) + \frac{U_j(S')}{C_0}\beta_j \\
&= (1-\alpha_j)\left(\frac{\sum_{i\in S}\beta_i U_i(S')}{\sum_{k\in S} U_k(S')}\right) + \alpha_j\beta_j. \qquad (21)
\end{aligned}$$

##### 3.2. Emergence of Global Epistasis

The term in the parentheses in Eq. (21) represents the internal allocation of the background species $S$ when species $j$ is present. Assuming the addition of species $j$ does not substantially change the internal allocation among species in $S$ ($\langle\beta\rangle_{S',\text{internal}} \approx \langle\beta\rangle_S$), we obtain:

$$\langle\beta\rangle_{S'} \approx (1-\alpha_j)\langle\beta\rangle_S + \alpha_j\beta_j. \tag{22}$$

Calculating the observed change $\Delta F$:

$$\begin{aligned}
F' - F &= C_0\left(\langle\beta\rangle_{S'} - \langle\beta\rangle_S\right) \\
&\approx C_0\left[(1-\alpha_j)\langle\beta\rangle_S + \alpha_j\beta_j - \langle\beta\rangle_S\right] \\
&\approx C_0\alpha_j\beta_j - C_0\alpha_j\langle\beta\rangle_S. \qquad (23)
\end{aligned}$$

Since $F = C_0\langle\beta\rangle_S$, we arrive at the linear relationship:

$$F' - F \approx \alpha_j(\beta_j C_0 - F). \tag{24}$$

#### CROSS-FEEDING MODEL

We extend the single-resource competition model to include a metabolic cascade. The community now contains two guilds: *producers* (biomass $x_p$) that consume a primary carbon source $C$, and *consumers* (biomass $x_k$) that consume a secondary metabolite $C_{cf}$ leaked by the producers. A fraction $p_p$ of each unit of $C$ taken up by producers is excreted as $C_{cf}$. Both guilds produce the byproduct $B$ that defines community function. Each producer is parameterized by its maximum growth rate $\mu_p$, biomass yield $Y_p$, and byproduct stoichiometry $\beta_p$; consumers analogously by $\mu_k$, $Y_k$, $\beta_k$. The two guilds use distinct Monod constants $K_M$ (on $C$) and $K_M^{cf}$ (on $C_{cf}$). The full dynamics are:

$$\begin{aligned}
\frac{dx_p}{dt} &= \frac{\mu_p C}{K_M + C}x_p \\
\frac{dx_k}{dt} &= \frac{\mu_k C_{cf}}{K_M^{cf} + C_{cf}}x_k \\
\frac{dC}{dt} &= -\sum_p \frac{1}{Y_p}\frac{\mu_p C}{K_M + C}x_p \\
\frac{dC_{cf}}{dt} &= \sum_p p_p\left(\frac{1}{Y_p}\frac{\mu_p C}{K_M + C}x_p\right) - \sum_k \frac{1}{Y_k}\frac{\mu_k C_{cf}}{K_M^{cf} + C_{cf}}x_k \\
\frac{dB}{dt} &= \sum_p \frac{\beta_p}{Y_p}\frac{\mu_p C}{K_M + C}x_p + \sum_k \frac{\beta_k}{Y_k}\frac{\mu_k C_{cf}}{K_M^{cf} + C_{cf}}x_k
\end{aligned} \tag{25}$$

As in the competitive model, simulations are run as batch cultures and function is the accumulated $B$ at carbon depletion.

#### Density-weighted correlation coefficient

To quantify the degree of linearity in a global epistasis plot we use a density-weighted Pearson correlation coefficient $\rho_w$ between the background function $F$ and the function gain $\Delta F = F' - F$. Across $2^{N-1}$ background communities, the empirical distribution of $F$ is highly non-uniform: many background compositions yield similar values of $F$, producing dense clusters along the $F$ axis (for instance near $F = 0$ when most low-producing backgrounds collapse onto each other), while other regions of $F$ are sampled by relatively few backgrounds. An unweighted Pearson correlation in this setting is dominated by the dense clusters along the $F$ axis. It can therefore miss a linear trend that spans the full range of $F$, or report spurious linearity when those clusters happen to lie along a line.

To correct for this, each point is weighted by the inverse of the marginal density of $F$, estimated from the data by a Gaussian kernel density estimate (KDE):

$$w_i \;=\; \frac{1}{\widehat{p}(F_i)}, \tag{26}$$

where $\widehat{p}(F)$ is the KDE of the empirical distribution of $\{F_i\}$. Points in sparse regions of $F$ are upweighted and points in dense regions downweighted, so the resulting correlation reflects linearity across the full range of $F$ rather than within the dominant cluster. Before computing weights we trim the top and bottom 1% of $\Delta F$ values to suppress outliers that arise from numerically extinct or near-empty backgrounds.

The density-weighted correlation is then the standard weighted Pearson formula:

$$\rho_w \;=\; \frac{\sum_i w_i(F_i - \bar{F}_w)(\Delta F_i - \overline{\Delta F}_w)}{\sqrt{\sum_i w_i(F_i - \bar{F}_w)^2 \, \sum_i w_i(\Delta F_i - \overline{\Delta F}_w)^2}}, \tag{27}$$

with weighted means $\bar{F}_w = \sum_i w_i F_i / \sum_i w_i$ and $\overline{\Delta F}_w$ defined analogously. With this definition $\rho_w \to -1$ when $\Delta F$ depends linearly and monotonically on $F$ across the full sampled range, while $\rho_w \to 0$ signals curvature or saturation that is invisible to a correlation dominated by the densely sampled cluster.

#### BREAKDOWN OF GLOBAL EPISTASIS

##### Niche partitioning

The model simulates a batch culture of $N$ species partitioned into two non-interacting guilds, $\mathcal{G}_A$ and $\mathcal{G}_B$, competing for two distinct resources, $C_A$ and $C_B$. The biomass $x_i$ of species $i$ accumulates via Monod growth on its guild-specific resource:

$$\frac{dx_{i\in\mathcal{G}_A}}{dt} = \mu_i \frac{C_A}{K_M + C_A} x_i, \quad \frac{dx_{i\in\mathcal{G}_B}}{dt} = \mu_i \frac{C_B}{K_M + C_B} x_i \tag{28}$$

Resource depletion is governed by the total uptake within each guild:

$$\frac{dC_A}{dt} = -\sum_{i\in\mathcal{G}_A} \frac{1}{Y_i}\frac{dx_i}{dt}, \quad \frac{dC_B}{dt} = -\sum_{i\in\mathcal{G}_B} \frac{1}{Y_i}\frac{dx_i}{dt} \tag{29}$$

The global community function $B$ represents the total integrated metabolic byproduct yield across both guilds:

$$\frac{dB}{dt} = \sum_{i=1}^{N} \beta_i \frac{1}{Y_i}\frac{dx_i}{dt} \tag{30}$$

The marginal contribution of a focal species, $\Delta F = B_{with} - B_{without}$, depends only on the occupancy of its own niche. Consequently, the global epistasis plot bifurcates: backgrounds consisting solely of the opposing group result in a zero-slope regime (independence), while backgrounds containing the same guild result in a negative-slope regime (competition).

##### Facilitation

The biomass $x_i$ of each species $i$ follows Monod growth kinetics, defined as:

$$\frac{dx_i}{dt} = \mu_i \frac{C}{K_M + C} x_i \tag{31}$$

where the primary resource $C$ is depleted by the total community uptake:

$$\frac{dC}{dt} = -\sum_{i=1}^{N} \frac{1}{Y_i}\frac{dx_i}{dt} \tag{32}$$

The total community function $B$ is the cumulative byproduct yield, governed by:

$$\frac{dB}{dt} = \sum_{i=1}^{N} \beta_i^{eff} \frac{1}{Y_i}\frac{dx_i}{dt} \tag{33}$$

Synergy is introduced via a conditional yield coefficient $\beta_i^{eff}$. If the facilitator (species $j = 0$) is present in the inoculum, the efficiency of all other species $i \neq 0$ is scaled by a synergy boost $\eta$:

$$\beta_{i\neq 0}^{eff} = \begin{cases} \beta_i(1+\eta) & \text{if } x_0(t) > \epsilon \\ \beta_i & \text{if } x_0(t) \le \epsilon \end{cases} \tag{34}$$

This formulation ensures that the marginal fitness contribution $\Delta F$ of species 0 is proportional to the total resource processed by the background species, leading to a positive slope in global epistasis scans.

#### SUPPLEMENTARY FIGURES

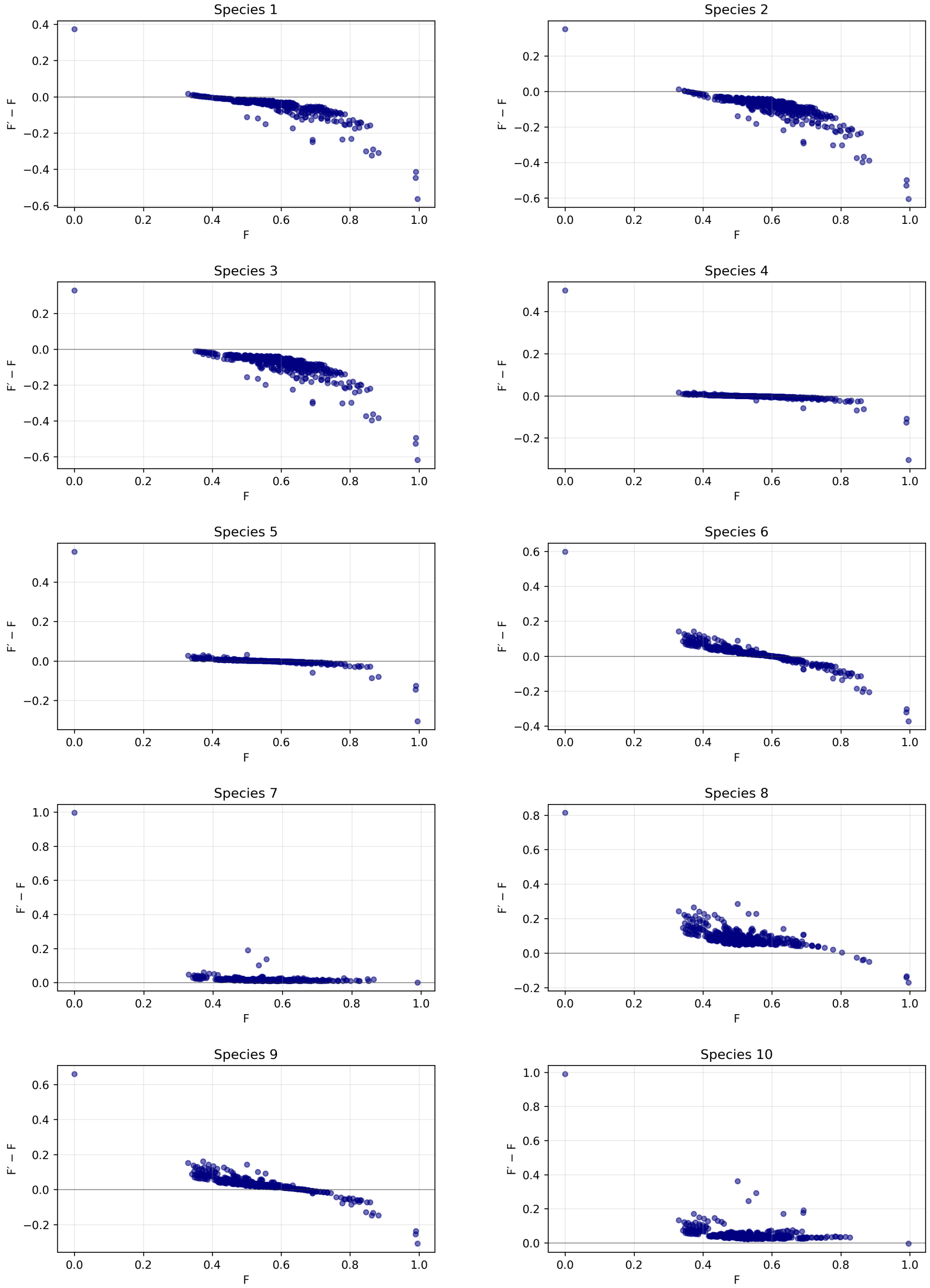


FIG. S1. **Global epistasis for all species in the community used for Figure 3**. The dashed lines are the analytical approximation to the slope given by equation 7 in the main text. Note the point at $F = 0$ is included here but omitted in the main text panels and corresponds to the target species being added to the empty community. The three example species in Figure 3 are 4, 2 and 6 shown in panels b-d respectively.

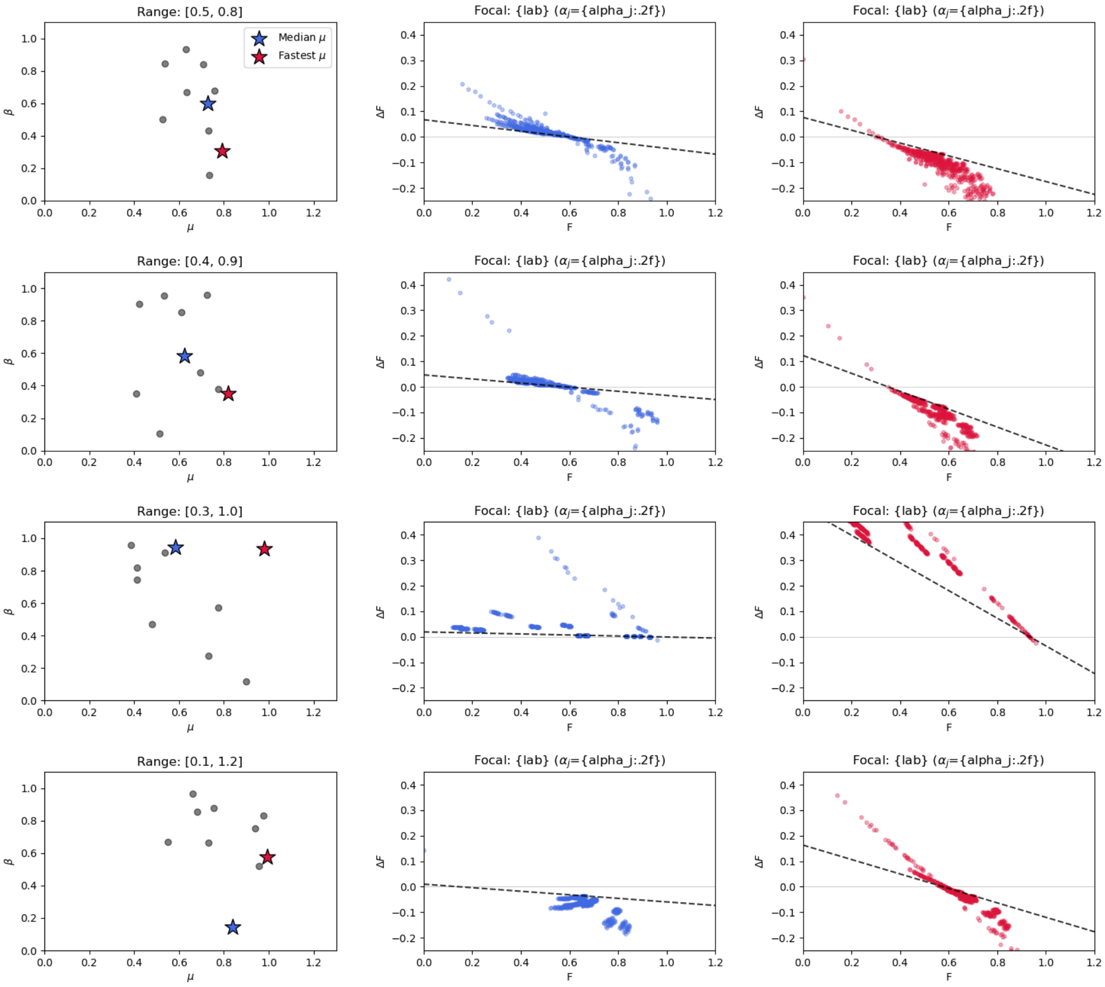


FIG. S2. **Stratification and branching of global epistasis under varying growth rate distributions for 10 species communities.** (Left Column) Phenotype maps showing the distribution of growth rates ($\mu$) and byproduct efficiencies ($\beta$) for $N = 10$ species. Red and blue stars indicate the 'Fastest $\mu$' and 'Median $\mu$' focal species, respectively. (Center and Right Columns) Global epistasis (GE) plots for the selected focal species across all $2^{N-1} = 512$ possible backgrounds. **Model Parameters:** Simulations were performed of the model shown in Eqns. 456 with $C_0 = 1.0$, half-saturation constant $K_M = 0.01$, and initial seed biomass $X_0 = 10^{-4}$. Yields ($Y$) were sampled from a lognormal distribution with a 20% coefficient of variation ($\sigma \approx 0.198$). **Dynamics:** From top to bottom, the range of $\mu$ is widened from a tight regime $[0.5, 0.8]$ to an extreme regime $[0.1, 1.2]$. The dashed black lines represent the analytical prediction $\Delta F = \alpha_j(\beta_j C_0 - F)$, where the resource share $\alpha_j$ is determined by a full $N$-species simulation. **Observations:** As the growth rate range expands, the 'Fastest $\mu$' strain exhibits significant branching, where discrete parallel lineages emerge. These branches represent different competitive regimes where the focal strain either dominates or shares resources with other high-growth competitors, causing the effective $\alpha_j$ to deviate from the community average.

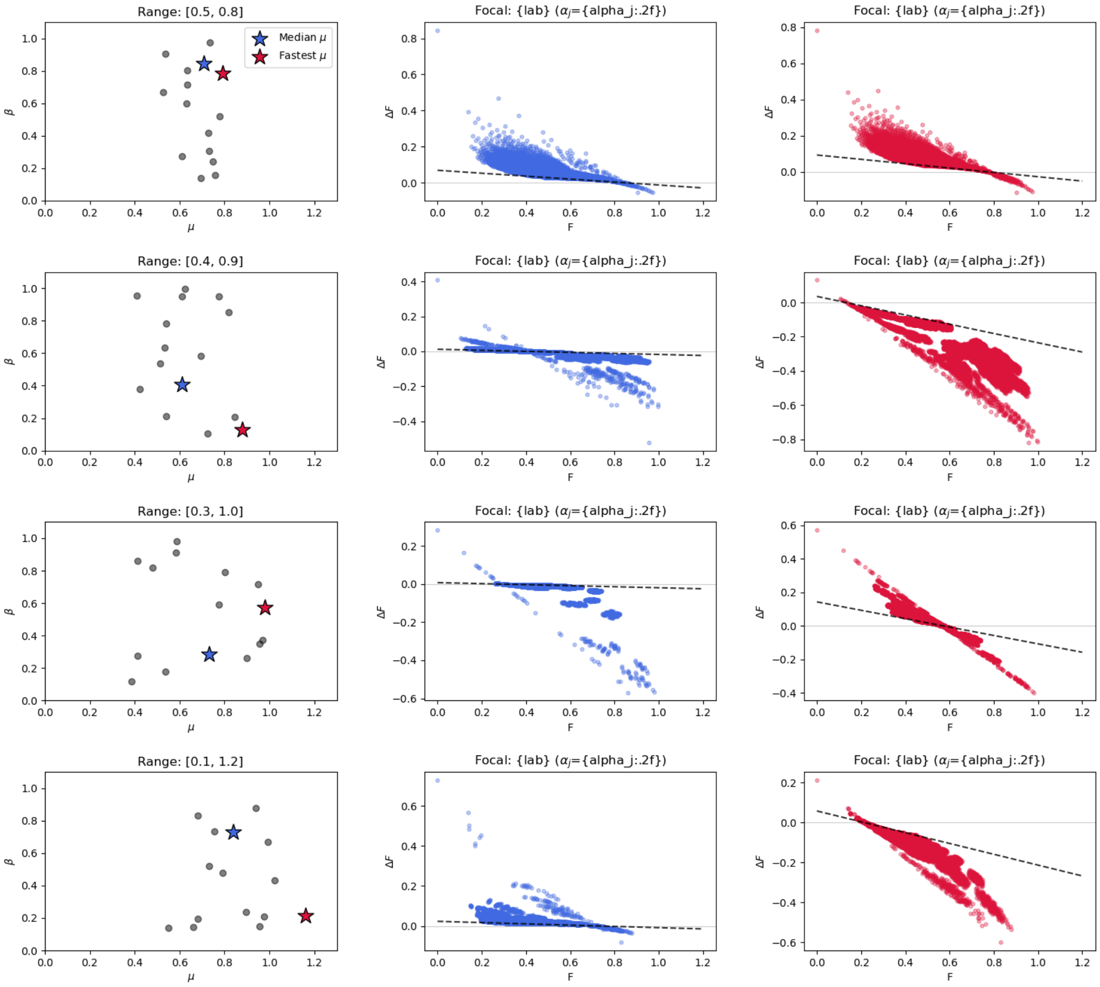


FIG. S3. **Stratification and branching of global epistasis under varying growth rate distributions for 15 species communities.** Identical to Figure S2 but for 15-species communities.

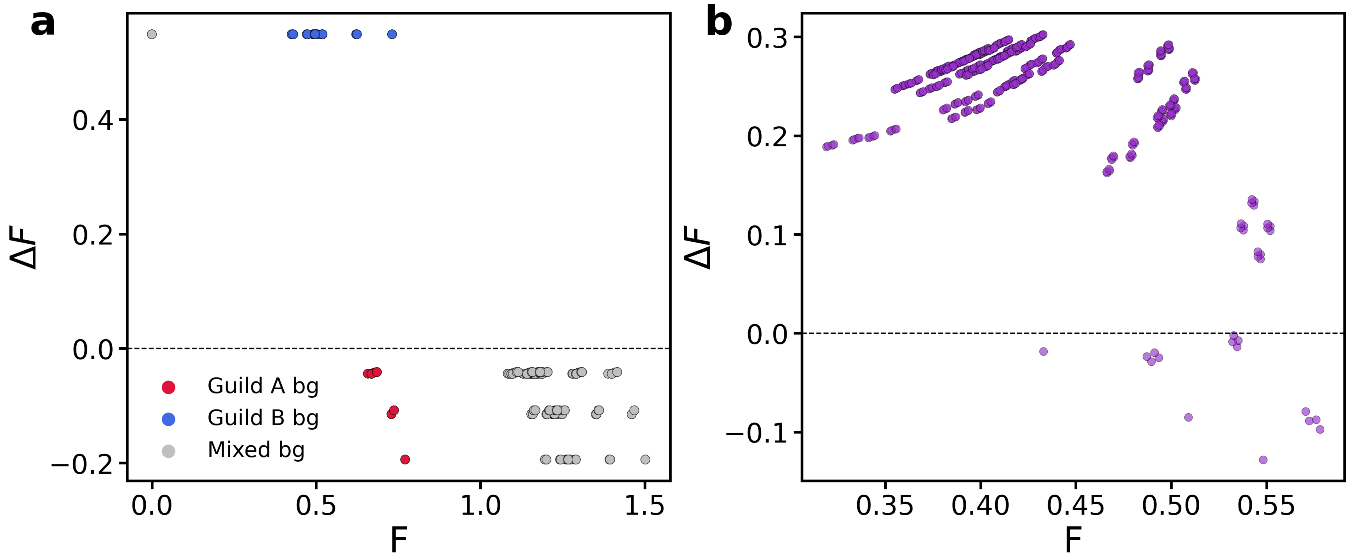


FIG. S4. **Niche partitioning and synergy break global epistasis patterns.** **(a)** Niche partitioning global epistasis plot for a single member of a 10-species community with two guilds that consume separate carbon sources. Points are colored by background composition (Guild A only, Guild B only, or mixed). **(b)** Global epistasis plot for a facilitator strain as described in the appendix. Note the positive slopes in various regions of the $\Delta F$ vs. $F$ plane.

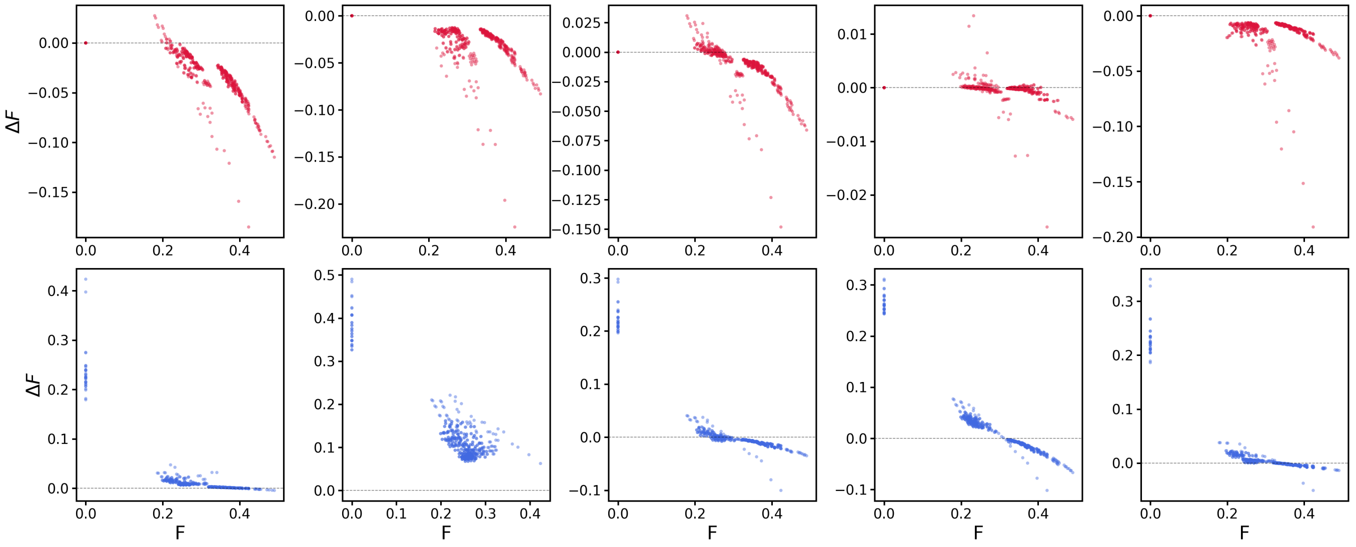


FIG. S5. **Strict cascade global epistasis.** GE plots for a community where the producers (strains degrading $C$ to $C_{cf}$ do not produce byproduct directly ($\beta_i = 0$ for $i \in P$). In this case the intercept for producers is always 0 since they produce no byproduct directly. Note in this case $p_i$ varies across strains unlike the simulations above where it is fixed across all producers.
